# Endosymbionts impact ladybird predation rates of aphids in a temperature-dependent manner

**DOI:** 10.1101/2024.09.20.614137

**Authors:** Katrine Bitsch, Perran A. Ross, Alex Gill, Qiong Yang, Monica Stelmach, Ashley Callahan, Michael Ørsted, Ary A. Hoffmann, Torsten N. Kristensen

## Abstract

Aphids are worldwide pests causing major economic losses to growers. Current management strategies rely heavily on pesticides, but some effective pesticides are being withdrawn and the efficiency of remaining pesticides is also decreasing as aphids build up resistance. Biological control using predators can provide a sustainable alternative to pesticides under some circumstances, while the deliberate introduction of facultative bacterial endosymbionts that induce host fitness costs and reduce plant virus transmission provides another potential future strategy to combat aphid pests. However, new control options should not be antagonistic, with the concern that the effectiveness of biocontrol might be altered by endosymbiont presence in hosts. We, therefore, tested if predation by two aphidophagous ladybirds, Adalia bipunctata and Harmonia conformis, on the green peach aphid, Myzus persicae, and the oat aphid, Rhopalosiphum padi, was affected by transinfected Rickettsiella viridis and both native and transinfected Regiella insecticola endosymbionts at different temperatures. The predation rate of aphids infected by either endosymbiont was higher at 14 ° C than the rate for uninfected aphids of both species, but the opposite pattern was apparent at 20 °C and for one host-endosymbiont combination at 26 °C. Overall, the results showed that higher temperatures increased predation, while differences between intermediate and low temperatures were species-dependent. No transmission of endosymbionts from aphid to ladybird through predation was detected. These findings point to a lack of consistent effects of the investigated endosymbionts on predation rates in these major aphid pests. The temperature dependence of endosymbiont-predation interactions suggests that the impact of seasonal climate should be considered when assessing the potential of endosymbionts in a biological control setting.

**HIGHLIGHTS:**

- Aphid endosymbiont effects on ladybird predation rates are temperature-dependent
- Ladybird predation rate of aphids increases as temperature rises
- Prey (aphid) endosymbionts are not transferred to predators (ladybirds)

## 1 INTRODUCTION

Aphids (Aphididae) represent economically important agricultural and horticultural pests globally, with damage pathways including phloem feeding, honeydew secretion, and the transfer of plant viruses (Jamin, 2023; Dedryver et al., 2010; Valenzuela and Hoffmann, 2015). To control aphids, chemical pesticides are typically applied but these can have severe negative consequences for biodiversity and ecosystem functioning (Geiger et al., 2010). Moreover, many aphids possess a remarkable ability to avoid or overcome the toxic effects of pesticides through rapid adaptation (Simon and Peccoud, 2018). Some aphids have developed resistance for many classes of insecticides, resulting in a requirement to increase dosages or adopt new pesticides (Bass et al., 2014). Alternative pest-control methods are therefore crucial for long-term sustainable control of aphids (Geiger et al., 2010), particularly when some effective chemicals are now being withdrawn due to environmental concerns (Pedersen and Nielsen 2017). One option is to increase the use of biological control, where natural biotic interactions are exploited to reduce pest infestations (Dedryver et al., 2010). In biological control, massreared natural enemies including predators are used to suppress pest populations, and/or other measures are undertaken to ensure that natural enemies are conserved in both greenhouse and field settings. Periodical releases of insect predators can be timed to match the level of pest infestation (van Lenteren et al., 2017).

Ladybirds (Coccinellidae) are some of the most used organisms in biological control of aphids (Symondson et al., 2002). However, the success of ladybirds for controlling aphids has been variable and dependent on numerous factors, including environmental conditions (van Lenteren et al., 2008; Sørensen et al., 2013; Uiterwaal and Delong, 2018). Many aspects of this strategy still need to explored to develop ladybirds as effective biological control agents of aphids (Symondson et al., 2002; van Lenteren et al., 2008; Jamin, 2023). One area where knowledge is limited is the interactions between ladybird predators and endosymbionts living inside the aphids (Dedryver et al., 2010).

Endosymbionts are microbes that live within with their host and can have effects ranging from parasitic to mutualistic (Henry et al., 2015, Hoffmann and Cooper, 2024, Zytynska et al., 2021). In aphids, the obligate endosymbiont Buchnera is typically seen as having a beneficial association with its host particularly by providing nutritional benefits (Douglas, 1998). Facultative endosymbionts can also provide benefits to hosts, which may be related to supplying essential nutrients but also include reduced susceptibility to parasitoids, parasites, predators, and pathogens (Losey et al., 1997; Łukasik et al., 2013; Oliver et al., 2003; Oliver et al., 2014). On the other hand, facultative endosymbionts can generate costs for hosts such as by decreasing host reproduction and reducing longevity (Laughton et al., 2014; Oliver et al., 2014; Vorburger and Gouskov, 2011) as well as increasing susceptibility to some natural enemies (Polin et al., 2014). These costs and benefits depend on the combination of the endosymbiont and host and may influence the field dynamics of endosymbionts in complex ways (Smith et al., 2021; Łukasik and Kolasa, 2024).

Facultative endosymbionts are now being used for suppressing populations of Aedes mosquitoes or replacing natural Aedes populations with individuals that suppress the transmission of dengue virus (Zheng et al., 2024; Utarnini et al., 2021; Nazni et al., 2019). These applications have sparked interest in utilizing endosymbionts in the control of agricultural pests including aphid species (Gu et al., 2023; Gong et al., 2020; Gou et al. 2023). Successful programs have relied on transinfections of endosymbionts from species naturally carrying the endosymbiont (donor) to species not known to carry the endosymbiont (recipient) resulting in the expression of novel traits in the recipient, including fitness costs and reduced viral transmission (Gong et al., 2020). Recently, transinfection of the endosymbiont Rickettsiella viridis into the green peach aphid, Myzus persicae, has been shown to induce several host fitness costs such as reduced fecundity and lowered heat tolerance, which could be utilized to inhibit the growth of M. persicae populations (Gu et al., 2023). Regiella insecticola is another candidate endosymbiont for pest management as it may influence plant virus transmission (Higashi et al., 2023) and has also now been transinfected into pest aphids (Gu et al., 2024).

Studies on the impact of native endosymbionts on predation by ladybirds have focused on the pea aphid, Acyrthosiphon pisum. Polin et al. (2014) found that pea aphids carrying the endosymbiont Hamiltonella defensa displayed reduced defensive behaviors against predators, including reduced rapid leg movements, and a reduced tendency to drop off surfaces and use their bodies to repel predators. Consequently, endosymbiont infections may make the host more vulnerable to attacks from ladybirds, potentially increasing the efficiency of biological control. This contrasts with studies showing that some native endosymbiont strains can decrease aphid susceptibility to parasitoids, at least in laboratory settings. This includes experiments showing that A. pisum is less susceptible to parasitoids when infected by H. defensa and that M. persicae is less susceptible to parasitoids when infected by some Re. insecticola strains (Oliver et al., 2008; Vorburger et al., 2010). However, there is little information on the interaction between cross-species transinfected endosymbionts and biocontrol except in the case of transinfected Ri. viridis which has no obvious impact on parasitism of M. persicae (Soleimannejad et al., 2023), although its impact on body colour might influence predation risk (Gu et al., 2023).

Beyond affecting host-predator interactions, endosymbionts may also benefit from natural enemies by facilitating their transfer between individuals or species. While aphid endosymbionts typically undergo vertical transmission from mother to offspring, horizontal transmission can also occur between individuals, especially if they share a food source (Sandström et al., 2001; Wilkinson et al., 2003; Russell et al., 2003; Gu et al., 2023; Soleimannejad et al., 2023) and particularly among conspecifics (Sanaei et al., 2021; Henry et al., 2013). In addition, horizontal transmission has been observed between predator and prey in a few cases, including those involving Wolbachia and Serratia symbiotica endosymbionts (Clec’h et al., 2013; Du et al., 2022). This could result in the spread of an introduced endosymbiont to other species in an environment which might be a concern in obtaining regulatory approval for deliberate releases.

Here we investigate effects of two endosymbionts, Ri. viridis and Re. insecticola, in the aphids M. persicae and Rhopalosiphum padi, on predation by two aphidophagous ladybirds, Adalia bipunctata and Harmonia conformis. The Ri. viridis infections considered here are transinfections, while the Re. insecticola infection is a native infection in M. persicae and transinfected in R. padi (Gu et al. 2024). We investigate predation at three temperatures (14°C, 20°C and 26°C) which is relevant because biological control should be efficient at a variety of ambient temperatures, as predation rates typically increase with increasing temperature, and because temperature influences endosymbiont density and distribution within a host (Gu et al. 2023; Uiterwaal and Delong, 2018; Sørensen et al., 2013). Further we tested if predation-mediated interspecies endosymbiont transfer between ladybirds and aphids harboring endosymbionts are observed. Results revealed highly temperature dependent effects of the endosymbionts and no transfer of endosymbionts between aphids and ladybird predators.

## 2 METHODOLOGY

### 2.1 Aphids

This study compared strains of aphids that were uninfected or infected with specific endosymbionts and shared the same genetic background. The Ri. viridis infected, M. persicae Rickettsiella +, and uninfected, M. persicae Rickettsiella -, strains of M. persicae were generated through microinjection from A. pisum that were naturally carrying Ri. viridis, as described in Gu et al. (2023). Aphid cultures of both M. persicae Rickettsiella + and M. persicae Rickettsiella - had been established from single females initially collected in Victoria, Australia. The same donor was used to generate R. viridis infected, R. padi Rickettsiella +, and uninfected, R. padi -, strains of R. padi as described in Chirgwin et al. (2023) and Gu et al. (2024). The native Re. insecticola infected, M. persicae Regiella +, and uninfected, M. persicae Regiella -, strains of M. persicae (cured and uncured strains 5.15) were described in Vorburger et al. (2010), with the M. persicae Regiella - strain produced by gentamicin treatment and the infection originating from a collection also made in Victoria, Australia. A different Regiella infected strain of M. persicae collected from Melbourne was used as the donor material for transinfected R. padi, R. padi Regiella +, produced through microinjection (Gu et al., 2024). The R. padi strain, R. padi -, used for transinfection of Regiella was the same as the one used for the Rickettsiella transinfection.

The M. persicae experiments were performed at Aalborg University, Denmark, where cultures were maintained on pak choi, (Brassica rapa subsp. chinesis), at 11 °C and under a 16:8 light:dark photoperiod. The R. padi experiments were performed at the University of Melbourne where cultures were maintained on wheat (Triticum aestivum var. Trojan) at 20 °C. Before performing experiments, populations of M. persicae were cultured with pak choi leaves in Petri dishes for two weeks under constant temperature conditions (20 °C) and a 12:12 light:dark photoperiod. For R. padi, populations were expanded on live wheat host plants for three weeks under constant temperature conditions (20 °C) and a 16:8 light:dark photoperiod in bugdorms (W24.5 x D24.5 x H63cm, 160 µm aperture, Australian Entomological Supplies Pty Ltd).

### 2.2 Ladybirds

This study used commercial stocks of two-spotted ladybirds, Adelia bipunctata, for the predation experiments with M. persicae, and large spotted ladybirds, Harmonia conformis, for predation on oat aphids. A. bipunctata was obtained from the Danish production facility Borregaard Bioplant ApS (http://uk.bioplant.dk), with received specimens being at both larval and egg stages. Large spotted ladybird was acquired from the Australian production facility Bugs for Bugs Pty Ltd (https://www.bugsforbugs.com.au), with received specimens being at the egg stage. All ladybirds developed on sterilized moth eggs. Following arrival, the ladybirds were kept at 20 °C with a 12:12 light:dark photoperiod.

### 2.3 Predation assay

To investigate ladybird predation rates of aphids at different temperatures, experiments were performed at 14 °C, 20 °C, and 26 °C. To ensure the aphids had the same age, an age-matching procedure was performed, whereby adult aphids (infected or uninfected from the same species/strain) were placed in Petri dishes (Ø60 mm) with 1% agar and host plant leaves for 24 hours at 20 °C. Subsequently, the adults were removed, while offspring stayed in the Petri dishes and matured undisturbed for an additional 72 hours before the experiments were initiated.

Thirty Petri dishes (Ø35 mm) were prepared for each endosymbiont status group (infected or uninfected strains of a particular species/endosymbiont combination). Each Petri dish contained a 1% agar solution and plant material for the aphids. For M. persicae, one cutout pak choi leaf disc (Ø20 mm) was used, while for R. padi there were three wheat leaves cut into 30 mm segments at the widest part of the leaf. Each Petri dish was labelled, and 15 age-matched aphids from one of the species/endosymbiont combinations was introduced. All Petri dishes were then transferred to a climate chamber held at the experimental temperature of 14 °C, 20 °C or 26 °C under a 12:12 light:dark photoperiod. After 24 hours, a ladybird larva, starved for 24 hours and weighed on an analytical balance (to the nearest 0.01 mg), was introduced into each Petri dish. For 36 hours, the number of remaining aphids in each dish was counted hourly. After the 36-hour period, counting was continued every 6 hours until all 15 aphids were consumed.

### 2.4 Endosymbiont detection in ladybirds

To investigate the potential transfer of the endosymbionts Re. insecticola and Ri. viridis from R. padi to large spotted ladybird (this work was not done for M. persicae), 68 ladybirds from the predation assay performed at 20 °C were tested. The ladybird larvae had consumed aphids infected with Re. insecticola, Ri. viridis or uninfected aphids. Half of the ladybird larvae from each group were starved for 24 hours after which they were stored individually in absolute ethanol in Eppendorf tubes at -80 ^°^C. The other half were also starved for 24 hours, then kept at 20 ^°^ C and fed uninfected R. padi until they reached the adult stage. They were then also stored individually in absolute ethanol in Eppendorf tubes at -80 ^°^ C. Based on the endosymbiont status of the consumed aphids and ladybird developmental stage, we tested the following numbers of ladybirds: for ladybird larvae, there were 12 ladybirds exposed to Re. insecticola, 15 exposed to Ri. viridis, and 12 uninfected; for adult ladybirds, there were 10 ladybirds exposed to Re. insecticola, 10 to Ri. viridis, and 9 uninfected.

To extract DNA from the ladybird specimens, each individual specimen was dried on a Kimwipe and put in a 1.7 mL Eppendorf tube with 3 µL of Proteinase-K (20 mg/ mL) solution, 150 µL 5% Chelex® (Bio-Rad Laboratories, Gladesville, NSW, Australia) solution and two metal beads (2 × 3 mm). The Eppendorf tubes were then grounded using a Qiagen TissueLyser II® at 25 Hertz for 4 minutes and the exoskeleton was removed after being centrifuged at 14,000 rpm for 1 min. The samples were then incubated at 65 ^°^C for one hour and then at 90 ^°^C for ten minutes. Lastly, the genomic DNA was diluted to a ratio of 1:10 with molecular-grade water and stored at -20^°^C.

### 2.5 Reference gene for H. conformis

A reference gene for Large spotted ladybird is essential, to confirm the concentration of ladybird DNA and quantify the potential endosymbionts. The designated primers were based on the best-performing reference genes investigated in relevant literatures (Yang et al., 2018; Lü et al., 2019; Qu et al., 2018) (Table 1). The studies referenced in Table 1 identified reference genes for the Multicoloured Asian ladybird, Harmonia axyridis and the 28-spotted potato ladybird, Henosepilachna vigintioctopunctata, both within the Coccinellidae. For each candidate reference gene (Table 1), conventional PCR assays were conducted at the annealing temperature recommended (Yang et al., 2018; Lü et al., 2019; Qu et al., 2018), and electrophoresis with post-staining of the gel using SYBR™ Safe (Invitrogen, Carlsbad, CA) was carried out with three samples from each developmental stage of the ladybirds and each endosymbiont group. Based on PCR, the annealing temperature of the two best-performing reference genes was adjusted to 58 °C, and PCR was rerun to confirm performance. The PCR conditions for DNA amplification began with a 10-min pre-incubation at 95 °C, followed by 40 cycles of 95 °C for 5 s, 58 °C for 15 s, and 72 °C for 30 s. A real-time quantitative PCR (RT-qPCR) was performed for the two reference genes using the same program and primers with conventional PCR, at the adjusted annealing temperature. The best-performing candidate, 28S ribosomal RNA (Yang et al. 2018), was selected as the reference gene (Figure 1).

**Table 1:**
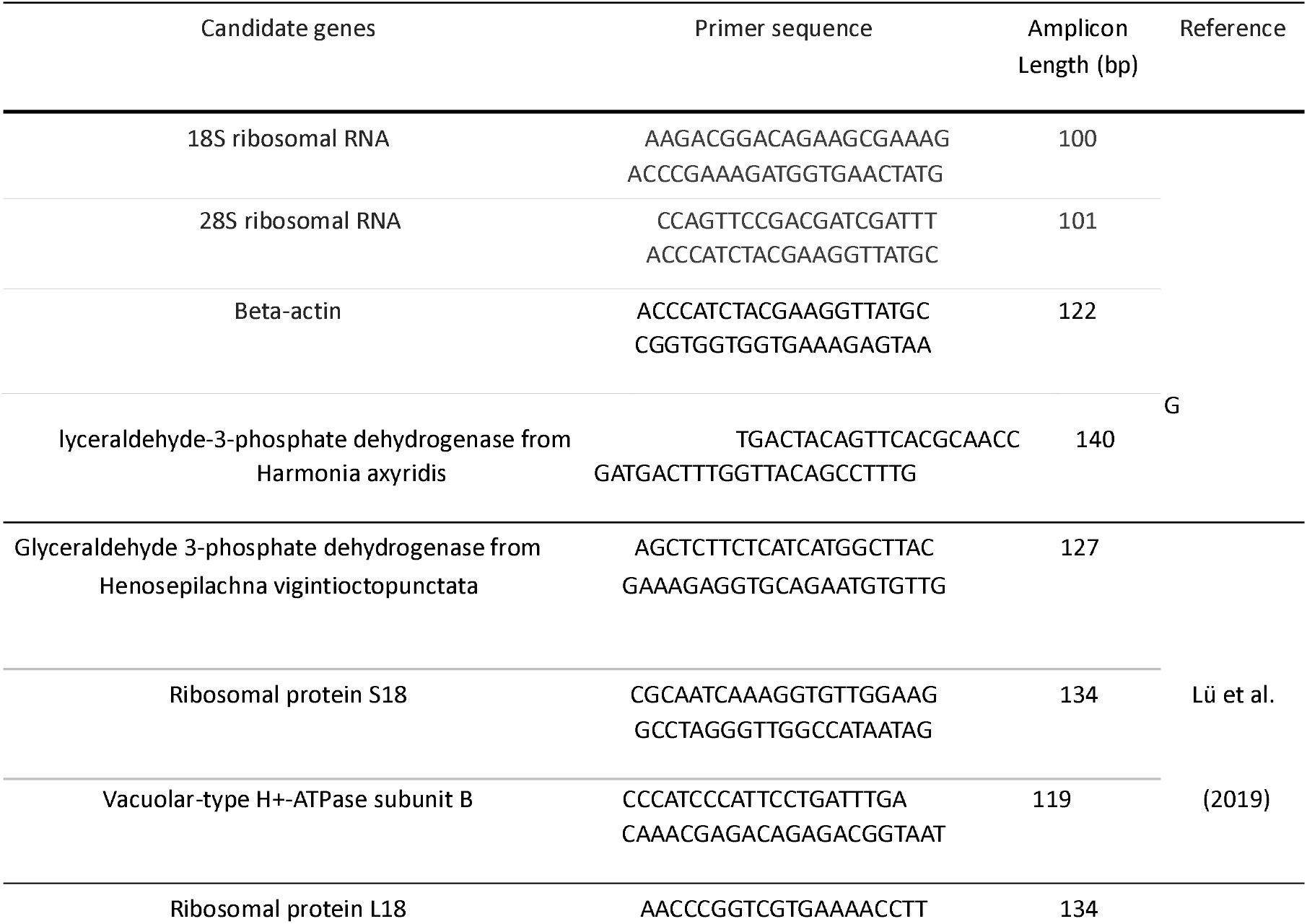
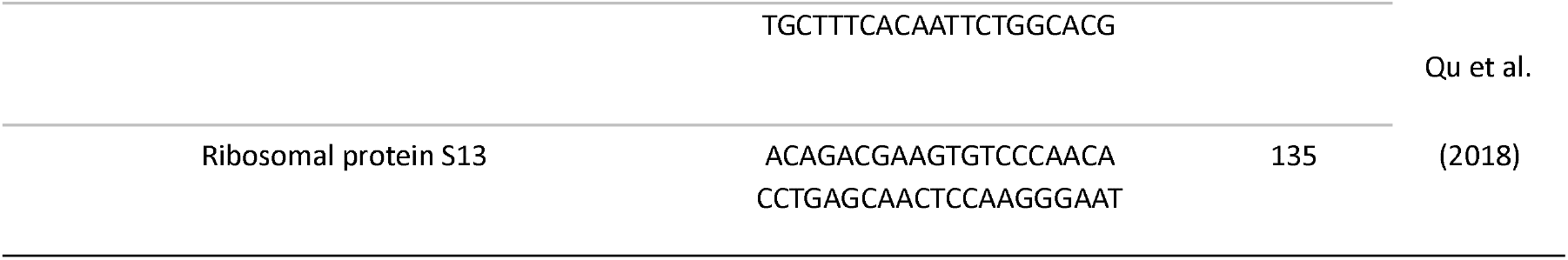
A list of candidate reference genes for the large spotted ladybird, along with their respective primer sequences, amplicon lengths (in base pairs), and references providing detailed information.

### 2.6 Detection of selected endosymbionts in ladybirds

To investigate the potential presence of the three known ladybird endosymbionts (Rickettsia spp., Spiroplasma spp. and Wolbachia spp.) and transfer of the two aphid endosymbionts (Ri. viridis and Re. insecticola), an RT-qPCR assay was performed. For the detection of each target endosymbiont, the designated primers from Table 2 were used, as well as the reference 28S ribosomal RNA gene (Yang et al., 2018). Crossing point (Cp) values of two technical replicate runs were averaged to produce the endosymbiont density when melting temperature (Tm) values were within the range of positive controls. However, there was inconsistency in the Tm values of Re. insecticola which led to further tests to confirm the presence/absence of this endosymbiont. Conventional PCR was performed on six suspected positive samples, six suspected negative samples, two positive controls, and two negative controls. After electrophoresis, DNA bands from four of the suspected positive samples and two of the expected negative samples were cut from the gel and isolated for purification before being sent for Sanger sequencing (Macrogen, Inc., Geumcheongu, Seoul, South Korea). Sequencing chromatograms were examined and processed with Geneious 9.18 software (Biomatters, Inc., Auckland, New Zealand).

**Table 2:**
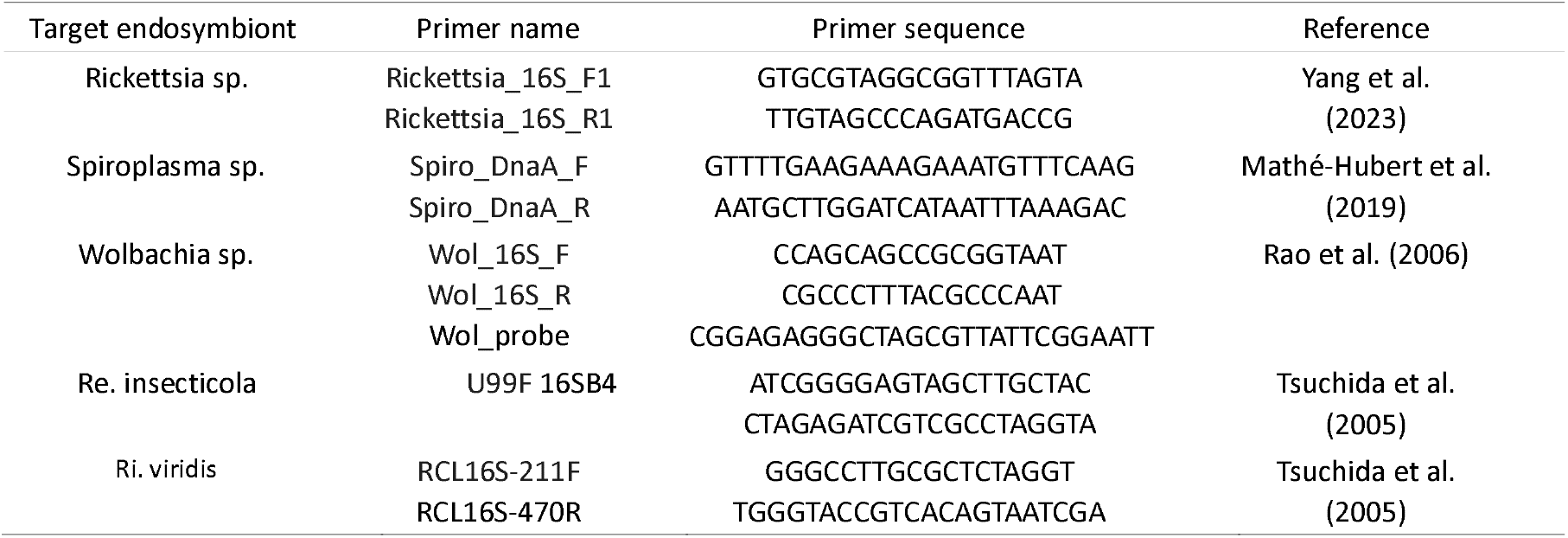
Primers used in this study to detect endosymbionts in the large spotted ladybird, along with their respective primer sequences and references providing detailed information.

**Table 3:** Log-rank (Mantel-Cox) test for significance, evaluating the difference between uninfected aphids and aphids infected with an endosymbiont (Ri. viridis or Re. insecticola).

| Comparison | 14°C |  |  | 20°C |  |  | 26°C |  |  |
| --- | --- | --- | --- | --- | --- | --- | --- | --- | --- |
| | $\chi^2$ | df | P | $\chi^2$ | df | P | $\chi^2$ | df | P |
| Large spotted ladybird |  |  |  |  |  |  |  |  |  |
| *R. padi* *Rickettsiella* + : - | 5.206 | 1 | 0.0225 | 11.44 | 1 | 0.0007 | 0.467 | 1 | 0.4670 |
| R. padi Regiella + : - | 25.26 | 1 | <0.0001 | 12.36 | 1 | 0.0004 | 3.009 | 1 | 0.0828 |
| Two-spotted ladybirds |  |  |  |  |  |  |  |  |  |
| M. persicae Rickettsiella + : - | 76.15 | 1 | <0.0001 | 12.54 | 1 | 0.0004 | 0.049 | 1 | 0.8250 |
| M. persicae Regiella + : - | 8.366 | 1 | 0.0038 | 25.35 | 1 | <0.0001 | 4.522 | 1 | 0.0335 |

### 2.7 Statistical Analysis

The effects of temperature and endosymbiont status on ladybird predation rates were analyzed with Kaplan-Meier survival curves and Wilcoxon rank-sum tests (Davidson-Pilon, 2024; Seabold and Perktold, 2010). Hazard ratios (Mantel-Haenszel) were calculated and used to assess the predation risk between uninfected and infected aphids across temperatures. If the 95% confidence intervals of the Hazard ratios did not include 1, the predation risk difference between infected and uninfected aphids was considered statistically significant. All statistical analyses were carried out in Python (Python 3.9).

## 3 RESULTS

### 3.1 Predation at different temperatures

Figure 2 illustrates the overall patterns of predation rates on aphids with and without endosymbiont infections across three temperature conditions (14 °C, 20 °C, and 26 °C). Generally, aphids infected with endosymbionts experience higher predation rates at 14 °C, lower rates at 20 °C, and similar rates at 26 °C compared to their non-infected counterparts. Large spotted ladybirds consumed R. padi infected with both endosymbionts significantly slower at 14 °C and significantly faster at 20 °C (Table 4) compared to the uninfected ones. There was no significant effect of endosymbiont infections at 26 °C for H. conformis. The same predation pattern at 14 °C and 20 °C was detected for A. bipunctata predating M. persicae. In addition, A. bipunctata consumed Re. insecticola infected M. persicae significantly faster than the uninfected ones at 26 °C.

**Table 4:** Hazard Ratio (Mantel-Haenszel) evaluating the overall difference between uninfected aphids and aphids infected with an endosymbiont (Ri. viridis or Re. insecticola). Significant differences are denoted by * (based on non-overlapping 95% confidence intervals). Hazard Ratios above one suggests that the risk of uninfected aphids being eaten is higher than the risk of infected being eaten and vice versa for values below one.

| Comparison | 14°C | 20°C | 26°C |
| --- | --- | --- | --- |
| Large spotted ladybird |  |  |  |
| R. padi Rickettsiella +/- | 1.18<sup>*</sup> | 0.75<sup>*</sup> | 0.95<sup>NS</sup> |
| R. padi Regiella +/- | 1.44<sup>*</sup> | 0.74<sup>*</sup> | 0.88<sup>NS</sup> |
| Two-spotted ladybirds |  |  |  |
| M. persicae Rickettsiella +/- | 1.94<sup>*</sup> | 0.77<sup>*</sup> | 1.02<sup>NS</sup> |
| M. persicae Regiella +/- | 1.23<sup>*</sup> | 0.69<sup>*</sup> | 1.19<sup>*</sup> |

**Table 5:** The observed Tm values for the screened endosymbionts and the differences between the ladybird samples and positive controls.

| Endosymbiont | Sample $T^m$ | Positive Control $T^m$ |
| --- | --- | --- |
| *Ri. viridis* | 70.60-79.52 | 87.66-88.00 |
| *Re. insecticola* | 81.04-86.95 | 87.08-87.12 |
| *Rickettsia* sp. | 79.78-80.54 | 82.92-83.10 |
| *Spiroplasma* sp. | 78.28-88.45 | 86.07-86.16 |
| *Wolbachia* sp. | 75.81-82.48 | 82.39-82.53 |

**Figure 2:**
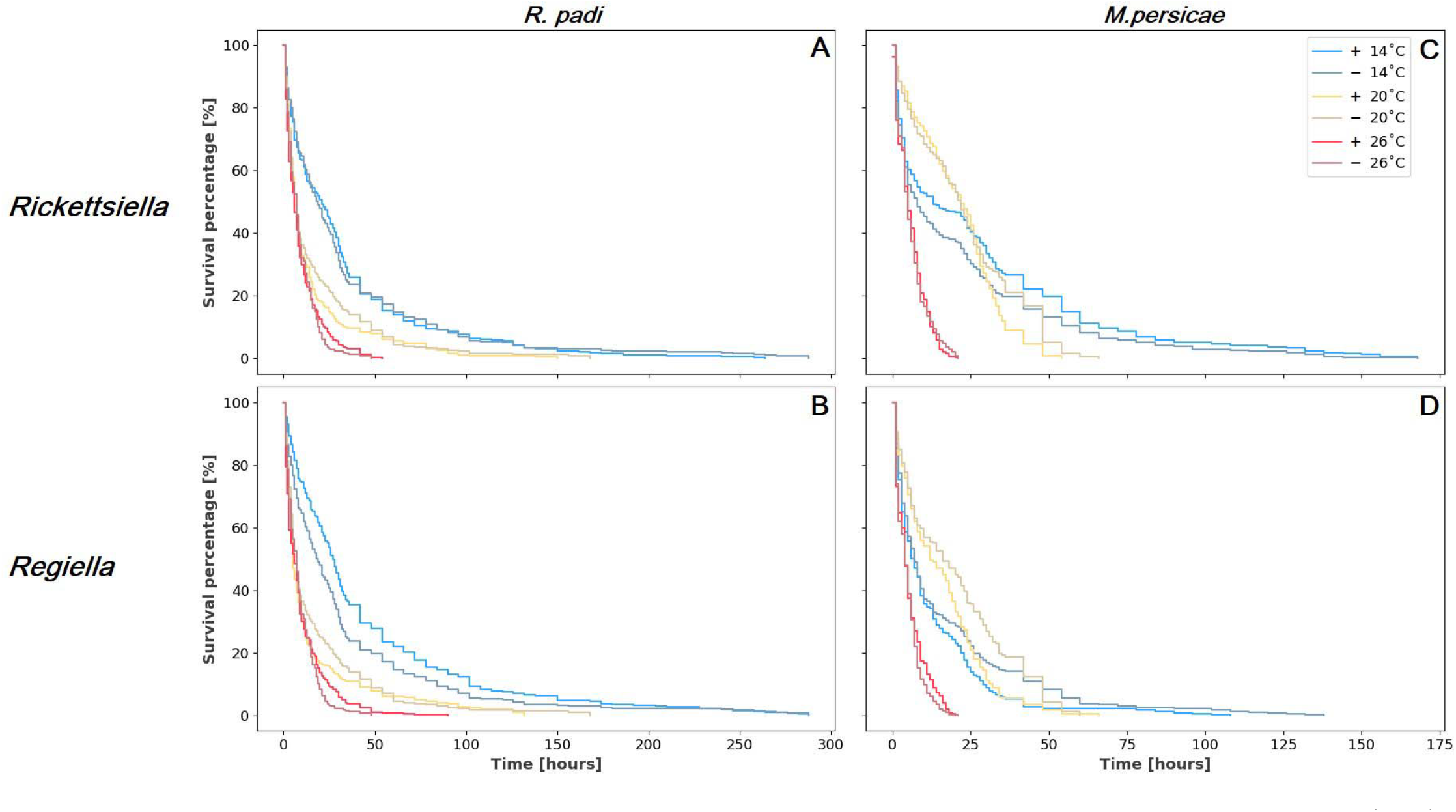
Kaplan-Meier survival curves of aphids with and without endosymbionts when exposed to ladybird predators at three temperatures (14 ^°^C, 20 ^°^C and 26 C). (A) and (B) is R. padi exposed to large spotted ladybird. (A) R.padi; Ri. viridis infected, +, and uninfected, -, reference. (B) R. padi; Re. insecticola infected, +, and unaffected, -. (C) and (D) is M. persicae exposed to two-spotted ladybird. (C) M. persicae; Ri. viridis infected, +, and uninfected, -, reference. (D) M. persicae; Re. insecticola infected, +, and unaffected, -, reference. Panels (A), (B), and (D) use the same line colour code as the legend of panel (C).

The Mantel-Haenszel Hazard Ratio (HR) revealed significant differences in the predation risk between uninfected and endosymbiont infected aphids, with the effect varying across temperatures and species (Table 4). At 14 °C, uninfected aphids had consistently higher predation risk compared to infected ones across all comparisons (e.g., R. padi with Rickettsiella [HR = 1.18, p < 0.05]; M. persicae with Rickettsiella [HR = 1.94, p < 0.05]). In contrast, at 20 °C, the risk was lower for uninfected aphids, as shown by Hazard Ratios below 1 (e.g., R. padi with Regiella [HR = 0.74, p < 0.05]). At 26 °C, most comparisons showed no significant difference, except for M. persicae with Regiella (HR = 1.19, p < 0.05).

Temperature significantly affected the feeding rate of ladybirds independent of species and endosymbiont infection status of the aphids (Figure 2 and Supplemental Table 2). Significant differences in ladybirds’ predation rate of aphids were observed across all temperatures, except for A. bipunctata preying on Re. insecticola -infected M. persicae, where no significant difference was found between 14 °C and 20 °C (Supplemental Table 2). This exception can be described by the Kaplan-Meier survival curves (Figure 2), where both A. bipunctata ladybirds infected with Re. insecticola and Ri. viridis, show a higher predation rate compared to at 20 °C during the beginning of the experiment. Furthermore, a significant difference was detected between the ladybirds predating R. padi, M. persicae Rickettsiella - and M. persicae Regiella (Supplementary Table 1).

### 3.2 No transfer of facultative endosymbionts through ladybird predation of aphids

The endosymbionts Ri. viridis and Re. insecticola were not present in the large spotted ladybird larvae stage nor when they developed into adult ladybirds. This finding is based on the fact that the qPCR screening for Rickettsiella of the ladybird samples did not indicate T^m^ values consistent with the positive controls which were screened concurrently. The qPCR screening for Regiella however did show T^m^ values similar to that of the positive controls (87.08-87.12), with some sample T^m^ values ranging from 85.08-86.95. To determine whether these were true Regiella detections, six samples were chosen which had relatively higher T^m^ values (85.08-86.95). Both positive controls and the six ladybird samples underwent a conventional PCR. Samples which showed bands at the same or similar size of the positive controls were excised from the gel along with the control sample bands, and the DNA purified using the Zymoclean™ Gel DNA Recovery Kit (Zymo Research). Sanger sequencing results showed that there was no Re. insecticola detected in the ladybird samples, while the positive controls were confirmed as Re. insecticola infection. Furthermore, no other native ladybird symbionts (Rickettsia sp., Spiroplasma sp., or Wolbachia sp.) were detected at either of the developmental stages.

## 4 DISCUSSION

### 4.1 Predation rates and implications for management strategies

Recent studies have pointed to the potential for using endosymbionts as a management strategy against aphids (Gu et al., 2023; Soleimannejad et al., 2023; Ross et al., 2024), which might constitute one sustainable alternative to pesticide use. However, it is important to investigate if endosymbiont manipulations can interfere with other pest management strategies such as using beneficial insects to control pests. Endosymbionts may impact on predator-prey interactions, and this led to our interest in exploring effects of aphid endosymbiont infection status on the rate of ladybird predation on these aphids and whether effects were temperature dependent. Further, we were interested in investigating whether endosymbionts introduced to aphids were detected in ladybirds eating them, which is of relevance from a regulatory perspective.

Across two study systems (A. bipunctata predating M. persicae and H. comformis predating A. padi), we show that the ladybird’s predation patterns are influenced by both temperature and the endosymbiont status of the aphid prey. However, the effect of endosymbionts on predation rate is strongly temperature dependent and depends on the interacting species involved (ladybirds and aphids). In a review of 23 ladybird species including A. bipunctata and Large spotted ladybird investigated here, Uiterwaal and Delong (2018), showed that ladybirds reduce pest populations more efficiently in warmer environments. Our study confirms this relationship in the case of M. persicae where the predation rate is highest at warm temperatures, 26 ^°^C. However, mixed results were found at intermediate, 20 ^°^C, and low, 14 ^°^ C, temperatures. M. persicae consumed faster at 14 ^°^C compared to at 20 ^°^C in the early phase of the experiment (Figure 2). This may depend on the temperatures aphids are kept at during maturation, where A. bipunctata were maintained at 5 ^°^C. The importance of acclimation for predation rates and predation efficiency were also emphasized by Sørensen et al. (2013).

The Hazard Ratio analyses reveal that endosymbiont infection significantly affects aphid predation risk, but effects are dependent on temperature and species (Table 4). At lower temperatures (14°C), uninfected aphids consistently had a higher risk of predation compared to infected ones, with hazard ratios ranging from 1.18 to 1.94. However, at higher temperatures, the protective effect of endosymbionts diminished, particularly at 26°C. These findings suggest that temperature modulates the impact of endosymbionts on aphid survival when exposed to predators, with the strongest effects observed at cooler temperatures. The temperature and species-specific effects of the endosymbionts are relevant to the implementation process of a pest management strategy involving ladybirds and endosymbionts. Regulatory approval is typically needed for endosymbiont release in natural aphid populations following similar procedures for the successful release of Wolbachia infected mosquitoes (Barro et al. 2011; Buchori et al., 2022). One component of this approval that also increases the effectiveness of the intervention is to take advantage of temperature dynamics revealed in our study, particularly for releases in temperate regions where aphid outbreaks are temperature dependent (Heie, 1994). In Denmark as an example, the peak season of M. persicae is from the start of summer until early fall (Miljøstyrelsen, 2017). In the midsummer months, the mean temperature closely resembles the experimental temperature of 20 ^°^C (Danmarks Meteorologiske Institut, 2023b). Based on our results, this implies higher predation rates on endosymbiont infected aphids, optimizing biocontrol effects. In contrast, during the early summer months, the mean temperature in Denmark is closer to the 14 ^°^C (Danmarks Meteorologiske Institut, 2023a) where the predation may be higher for the aphids without endosymbionts. Releases at this time could be utilised to facilitate the spread of the endosymbiont, prior to the period when aphid damage is at a maximum along with predation rates by ladybirds, helping in biocontrol.

Thus, tailoring release timing of endosymbiont infected aphids to specific thermal environmental conditions may aid the control of aphids while reducing the need of chemical pesticides. Such a strategy can be informed by results from predictive models assessing temporal and spatial distributions of outbreaks of certain aphids (or other pest species) (Damgaard et al., 2020). However, large scale greenhouse and open field experiments should be conducted to indicate whether these approaches are effective in an applied context.

### 4.2 Endosymbionts transmission and

Environmental impacts of releasing endosymbiont transinfections in aphids will need to be assessed before potentially deploying endosymbionts as a management strategy. A hazard addressed here is the potential horizontal transfer of the endosymbiont to other invertebrates across food webs. Horizontal and vertical transfer of Ri. viridis between aphids have already been detected by Gu et al. (2023) in M. persicae and the presence of this infection in an aphid parasitoid was noted by Soleimannejad et al. (2023). However, in parasitoids, no transfers were detected between parasitoid generations or into uninfected aphids (Yang et al., 2023). Nevertheless, other studies have identified inter-species transmissions of endosymbionts such as Serratia symbiotica and Wolbachia between prey and predators (Clec’h et al., 2013; Du et al., 2022) so we suspect that this issue needs to be considered on a case-by-case basis. In our study on ladybirds, we did not find evidence of the endosymbionts Ri. viridis and Re. insecticola being present in ladybird larvae or adults that had eaten infected aphids, suggesting inter-species transmission is not likely via this pathway.

In future research, several steps could be undertaken to advance our understanding and application of endosymbionts in aphid management. Computational models could be an important tool to predict the dynamics of outbreaks and endosymbiont-host interactions under various environmental conditions and thereby evaluate the potential impact of endosymbionts on aphid populations (Slavenko et al. 2024). To empirically validate the accuracy of the models, tests should be performed under more production relevant environmental conditions in greenhouses, to assess the effectiveness and stability of introducing endosymbionts. Following greenhouse trials, field releases should be carefully designed and implemented to test the real-world applicability and scalability of these interventions, with an emphasis on environmental safety and long-term sustainability. Additionally, the exploration of endosymbionts not investigated here with different or complementary functions could provide a broader toolkit for biological control, enhancing the resilience of pest management strategies against various aphid species and environmental stresses. These steps may collectively contribute to the development of innovative and eco-friendly approaches to aphid management and reduce reliance on chemical pesticides.

## 5 CONCLUSION

We found that the predation rate of aphids by ladybirds was affected by temperature, with increased predation rates at higher temperatures as also observed in other studies, but with mixed results at intermediate and low temperatures. The effects of aphid endosymbiont status on predation rates were also temperature dependent. Thus, our study highlights the complexity of interactions between temperature and endosymbiotic infection status of aphids, on ladybird predation efficiency. These findings emphasize the need to consider seasonal and geographic climate variation when incorporating such biocontrol elements into aphid management. We found no evidence that inter-species transmission occurred when Large spotted ladybird preyed on R. padi infected with the endosymbionts Ri. viridis and Re. insecticola, suggesting that predation does not lead to a risk of the endosymbionts spreading from R. padi to Large spotted ladybird and through this pathway to other species.

## 7 ACKNOWLEDGEMENT

We are grateful to Martin Kynde, Eloise Ansermin, Ashritha Prithiv Sivaji Dorai, and Katrine Wagner Thiessen for their academic advice and assistance with experimental tasks.

## 8 AUTHOR CONTRIBUTIONS

Katrine Bitsch Thomsen: Conceptualization, Formal analysis, Funding acquisition, Investigation, Methodology, Project administration, Visualization, Writing – original draft

Perran A. Ross: Conceptualization, Methodology, Supervision, Writing – review and editing

Alex Gill: Conceptualization, Methodology, Supervision, review and editing

Qiong Yang: Methodology, Formal analysis, Writing – review and editing

Monica Stelmach: Methodology, Writing – review and editing

Ashley Callahan: Methodology, Supervision

Michael Ørsted: Methodology, Supervision, Writing – review and editing

Ary A. Hoffmann: Conceptualization, Funding acquisition, Methodology, Supervision, Writing – review and editing

Torsten N. Kristensen: Conceptualization, Funding acquisition, Methodology, Supervision, Writing – review and editing

## REFERENCES

Barro, P., Murphy B., Jansen, C., and Murray, J. (2011). The proposed release of the yellow fever mosquito, Aedes aegypti containing a naturally occurring strain of Wolbachia pipientis, a question of regulatory responsibility. Journal für Verbraucherschutz und Lebensmittelsicherheit, 6.Suppl 1:33–40.

Bass, C., Puinean, A. M., Zimmer, C. T., Denholm, I., Field, L. M., Foster, S. P., Gutbrod, O., Nauen, R., Slater, R., and Williamson, M. S. (2014). The evolution of insecticide resistance in the peach potato aphid, myzus persicae. Insect biochemistry and molecular biology, 51:41–51.

Buchori, D., Mawan, A., Nurhayati, I., Aryati, A., Kusnanto, H., and Hadi, U. K. (2022). Risk assessment on the release of Wolbachia-infected Aedes aegypti in yogyakarta, indonesia. Insects, 13:924.

Chirgwin, E., Yang, Q., Umina, P. A., Thia, J. A., Gill, A., Song, W., Gu, X., Ross, P.A., Wei, S.-J., and Hoffmann, A. A. (2024). Barley yellow dwarf virus influences its vector’s endosymbionts but not its thermotolerance. Microorganisms, 12, 10.

Clec’h, W. L., Chevalier, F., Genty, L., Bertaux, J., Bouchon, D., and Sicard, M. (2013). Cannibalism and predation as paths for horizontal passage of Wolbachia between terrestrial isopods. Plos one, 8:e60232.

Damgaard, C., Bruus, M., Axelsen, J. A. (2020). The effect of spatial variation for predicting aphid outbreaks. Journal of applied entomology 144, 4:263–269

Danmarks Meteorologiske institute (2023a). Vejrog klimadata, Danmark, sæson oversigt 2023 - efterår. https://www.dmi.dk/fileadmin/user_upload/Afrapportering/Saesonoversigter/Oversigt_2023_efter%C3%A5r.pdf.

Danmarks Meteorologiske Institut (2023b). Vejr-og klimadata, Danmark, sæsonoversigt 2023 - sommer. https://www.dmi.dk/fileadmin/user_upload/Afrapportering/Saesonoversigter/Oversigt_2023_sommer.

Davidson-Pilon, C. (2024). Lifelines, survival analysis in Python. Zenodo, https://www.lifelines.readthedocs.io/en/latest/. Version v0.28.0, employed in Python version 3.8.

Dedryver, C.-A., Le Ralec, A., and Fabre, F. (2010). The conflicting relationships between aphids and men: A review of aphid damage and control strategies. Comptes rendus biologies, 333:539–553.

Douglas, A.E. (1998). Nutritional interactions in insect-microbial symbioses: aphids and their symbiotic bacteria Buchnera. Annual review of entomology, 43:17–37.

Du, X.-Y., Yang, H.-Y., Gong, S.-R., Zhang, P.-F., Chen, P.-T., Liang, Y.-S., Huang, Y.-H., Tang, X.-F., Chen, Q.-K., Clercq, P. D., Li, H.-S., and Pang, H. (2022). Aphidophagous ladybird beetles adapt to an aphid symbiont. Functional ecology, 36(10):2593– 2604.

Geiger, F., Bengtsson, J., Berendse, F., Weisser, W. W., Emmerson, M., Morales, M. B., Ceryngier, P., Liira, J., Tscharntke, T., Winqvist, C., Eggers, S., Bommarco, R., Pärt, T., Bretagnolle, V., Plantegenest, M., Clement, L. W., Dennis, C., Palmer, C., Oñate, J. J., Guerrero, I., Hawro, V., Aavik, T., Thies, C., Flohre, A., Hänke, S., Fischer, C., Goedhart, P. W., and Inchausti, P. (2010). Persistent negative effects of pesticides on biodiversity and biological control potential on european farmland. Basic and applied ecology, 11:97–105.

Gong, J., Li, Y., Li, T., Hong, X., Hoffmann, A., Xi, Z., Hu, L., Zhang, D., and Baton, L. (2020). Stable introduction of plant-virusinhibiting Wolbachia into planthoppers for rice protection. Current biology. 30:4837–4845.

Gu, X., Ross, P. A., Yang, Q., Gill, A., Umina, P. A., and Hoffmann, A. A. (2024). Genetic and environmental factors influence the success of endosymbiont transfers in pest aphids. Environmental microbiology, (in press)

Gu, X., Ross, P. A., Gill, A., Yang, Q., Ansermin, E., Sharma, S., Soleimannejad, S., Sharma, K., Callahan, A., Brown, C., Umina, P. A., Kristensen, T. N., and Hoffmann, A. A. (2023). A rapidly spreading deleterious aphid endosymbiont that uses horizontal as well as vertical transmission. Proceedings of the national academy of sciences, 120(18):e2217278120.

Guo, Y., Shao, J., Wu, Y., and Li, Y. (2023). Using Wolbachia to control rice planthopper populations: progress and challenges. Frontiers in microbiology, 14.

Heie, O. E. (1994). Why are there so few aphid species in the temperate areas of the southern hemisphere? European journal of entomology, 91(1):127–133.

Henry, L., Maiden, M., Ferrari, J., and Godfray, C. (2015). Insect life history and the evolution of bacterial mutualism. Ecology letters, 18.

Henry, L. M., Peccoud, J., Simon, J.-C., Hadfield, J. D., Maiden, M. J., Ferrari, J., and Godfray, C. J. (2013). Horizontally transmitted symbionts and host colonization of ecological niches. Current biology, 23(17):1713–1717.

Higashi, C. H. V., Nichols, W. L., Chevignon, G., Patel, V., Allison, S. E., Kim, K. L., Strand, M. R., and Oliver, K. M. (2023). An aphid symbiont confers protection against a specialized rna virus, another increases vulnerability to the same pathogen. Molecular biology, 32(4):936–950.

Hoffmann, A. A., and Cooper, B. S., (2024). Describing endosymbiont–host interactions within the parasitism–mutualism continuum. Ecology and evolution, 14:7.

Intergovernmental Panel on Climate Change (2023). Climate change 2023: Synthesis report, assessment report of the [core writing team, h. lee and j. romero (eds.)]. IPCC, pages 35–115.

Jamin, A. (2023). The peach potato aphid (Myzus persicae): Ecology and management, chapter 2.3. Number ISBN: 9781003400974 in 1. CRC Press.

Laughton, M. A., Fan, H. M., & Gerardo, M. N., (2014). The combined effects of bacterial symbionts and aging on life history traits in the pea aphid, Acyrthosiphon pisum. Applied and environmental microbiology, 80:470–477.

Leys, C., Ley, C., Klein, O., Bernard, P., and Licata, L. (2013). Detecting outliers: Do not use standard deviation around the mean, use absolute deviation around the median. Journal of experimental social psychology, 49:764–766.

Losey, J.E., Ives, A.R., Harmon, J., Ballantyne, F. & Brown, C. (1997) A polymorphism maintained by opposite patterns of parasitism and predation. Nature, 388:269–272.

Lü, J., Chen, S., Guo, M., Ye, C., Qiu, B., Wu, J., Yang, C., and Pan, H. (2019). Selection and validation of reference genes for rt-qpcr analysis of the ladybird beetle Henosepilachna vigintioctopunctata. Frontiers in physiology, 10.

łukasik, P., and Kolasa, M. R. (2024). With a little help from my friends: the roles of microbial symbionts in insect populations and communities. Philosophical transactions of the Royal Society of London. Series B, Biological sciences, 379.

łukasik, P., van Asch, M., Guo, H.F., Ferrari, J. & Godfray, H.C.J. (2013) Unrelated facultative endosymbionts protect aphids against a fungal pathogen. Ecology letters, 16:214–218.

Mathé-Hubert, H., Kaech, H., Ganesanandamoorthy, P., and Vorburger, C. (2019). Evolutionary costs and benefits of infection with diverse strains of Spiroplasma in pea aphids. Evolution, 73:1466–1481.

Miljøstyrelsen (2017). Udvikling og afprøvning af koncept for beslutningsstøttesystem for bekæmpelse af skadevoldere i landbruget. https://www2.mst.dk/Udgiv/publikationer/2017/09/978-87-93614-25-3.pdf.

Nazni, W. A., Hoffmann, A. A., NoorAfizah, A., Cheong, Y. L., Mancini, M. V., Golding, N., Kamarul, G. M. R., Arif, M. A. K., Thohir, H., NurSyamimi, H., ZatilAqmar, M. Z., NurRuqqayah, M., NorSyazwani, A., Faiz, A., Irfan, F. M. N., Rubaaini, S., Nuradila, N., Nizam, N. M. N., Irwan, S. M., Endersby-Harshman, N. M., and Sinkins, S. P. (2019). Establishment of Wolbachia strain wAlbB in Malaysian populations of Aedes aegypti for dengue control. Current biology. 24:4241–4248.

Oliver, K. M., Campos, J., Moran, N. A., and Hunter, M. S. (2008) Population dynamics of defensive symbionts in aphids. Proc. R. Soc. B. 275:293–299

Oliver, K. M., Russell, J. A., Moran, N. A. & Hunter, M. S. (2003) Facultative bacterial symbionts in aphids confer resistance to parasitic wasps. Proceedings of the national academy of sciences, 100:1803–1807.

Oliver, K. M., Smith, A. H., and Russell, J. A. (2014). Defensive symbiosis in the real world – advancing ecological studies of heritable, protective bacteria in aphids and beyond. Functional ecology, 28:341–355.

Pecenka, J., Ingwell, L., Foster, R., Krupke, C., and Kaplan, I. (2021). Ipm reduces insecticide applications by 95%. Proceedings of the national academy of sciences, 118.

Pedersen, A. B., and Nielsen H. Ø. (2017). Effectiveness of pesticide policies. In environmental pest management. eds M. Coll and E. Wajnberg, ch13:297–324

Polin, S., Gallic, J.-F. L., Simon, J.-C., Tsuchida, T., and Outreman, Y. (2015). Conditional reduction of predation risk associated with a facultative symbiont in an insect. PLoS ONE, 10(11):e0143728.

Polin, S., Simon, J.-C., and Outreman, Y. (2014). An ecological cost associated with protective symbionts of aphids. Ecology and evolution, 4:836–840.

Popovici, J., Moreira, L. A., Poinsignon, A., Iturbe-Ormaetxe, I., McNaughton, D., & O’Neill, S. L. (2010). Assessing key safety concerns of a Wolbachia-based strategy to control dengue transmission by Aedes mosquitoes. Memorias do instituto oswaldo cruz, 105:957–964.

Qu, C., Wang, R., Che, W., Zhu, X., Li, F., and Luo, C. (2018). Selection and evaluation of reference genes for expression analysis using quantitative real-time pcr in the asian ladybird Harmonia axyridis (coleoptera: Coccinellidae). Plos one, 13:1–15.

Rao, R. U., Atkinson, L. J., Ramzy, R. M. R., Helmy, H., Farid, H. A., Bockarie, M. J., Susapu, M., Laney, S. J., Williams, S. A., and Weil, G. J. (2006). A real-time pcr-based assay for detection of Wuchereria bancrofti dna in blood and mosquitoes. The American journal of tropical medicine and hygiene, 74:826–832.

Ross, P., Tyrilos, M., Durugkar, N., Gill, A., Jonge, N., Yang, Q., Gu, X., Hoffmann, A. A., and Kristensen, T. N. (2024). Deleterious effects of the endosymbiont Rickettsiella viridis in Myzus persicae are environmentally dependent, Journal of pest science, 1–14.

Russell, J. A., Latorre, A., Sabater-Muñoz, B., Andrés Moya, A., and Moran, N. (2003). Side-stepping secondary symbionts: Widespread horizontal transfer across and beyond the aphidoidea. Molecular ecology, 12(4):1061–1075.

Sanaei, E., Charlat, S., and Engelstädter, J. (2021). Wolbachia host shifts: Routes, mechanisms, constraints and evolutionary consequences. Biological reviews, 96:433–453.

Sandström, J. P., Russell, J. A., White, J. P., and Moran, N. A. (2001). Independent origins and horizontal transfer of bacterial symbionts of aphids. Molecular ecology, 10(1):217–228.

Seabold, S. and Perktold, J. (2010). Statsmodels: Econometric and statistical modelling with python. Proceedings of the 9th Python in Science Conference, https://www.statsmodels.org/stable/. employed in Python version 3.8.

Simon, S., and Peccoud J. (2018). Rapid evolution of aphid pests in agricultural environments. Current opinion in insect science, 26:17–24.

Slavenko, A., Ross, P. A., Mata, L., Hoffmann, A. A., and Umina, P. A. (2024). Modelling the spread of a novel endosymbiont infection in field populations of an aphid pest. Ecological Modelling. 497:110851.

Smith, A., O’Connor, M., Deal, B., Kotzer, C., Lee, A., Wagner, B., Joffe, J., Woloszynek, S., Oliver, K., and Russell, J. (2021). Does getting defensive get you anywhere?—Seasonal balancing selection, temperature, and parasitoids shape real-world, protective endosymbiont dynamics in the pea aphid. Molecular ecology. 30:2449–2472.

Soleimannejad, S., Ross, P. A., and Hoffmann, A. A. (2023). Effect of rickettsiella viridis endosymbionts introduced into Myzus persicae aphids on parasitism by Diaeretiella rapae: A combined strategy for aphid control. Biological control, 187:105377.

Symondson, W. O. C., Sunderland, K. D., and Greenstone, M. H. (2002). Can generalist predators be effective biocontrol agentŠ Annual review of entomology, 47:561–94.

Sørensen, C. H., Toft, S., and Kristensen, T. N. (2013). Cold-acclimation increases the predatory efficiency of the aphidophagous coccinellid Adalia bipunctata. Biological control, 65:87–94.

Tsuchida, T., Koga, R., Fujiwara, A., and Fukatsu, T. (2014). Phenotypic effect of “Candidatus Rickettsiella viridis,” a facultative symbiont of the pea aphid (Acyrthosiphon pisum), and its interaction with a coexisting symbiont. Applied and environmental microbiology, 80:525–533.

Tsuchida, T., Koga, R., Meng, X. Y., Matsumoto, T., and Fukatsu, T. (2005). Characterization of a facultative endosymbiotic bacterium of the pea aphid Acyrthosiphon pisum. Microbial ecology, 49:126–133.

Uiterwaal, S. and Delong, J. (2018). Multiple factors, including arena size, shape the functional responses of ladybird beetles. Journal of applied ecology, 55.

Utarini, A., Indriani, C., Ahmad, R. A., Tantowijoyo, W., Arguni, E., Ansari, M. R., Supriyati, E., Wardana, D. S., Meitika, Y., Ernesia, I., Nurhayati, I., Prabowo, E., Andari, B., Green, B. R., Hodgson, L., Cutcher, Z., Rancès, E., Ryan, P. A., O’Neill, S. L., Dufault, S. M., (2021). Efficacy of Wolbachia-infected mosquito deployments for the control of dengue. The New England journal of medicine, 23:2177–2186.

Valenzuela, I. and Hoffmann, A. A. (2015). Effects of aphid feeding and associated virus injury on grain crops in Australia. Austral entomology, 54:292–305.

van Lenteren, J., Bolckmans, K., Köhl, J., Ravensberg, W., and Urbaneja, A. (2017). Biological control using invertebrates and microorganisms: Plenty of new opportunities. Biocontrol, 63.

van Lenteren, J., Loomans, A., Babendreier, D., and Bigler, F. (2008). Harmonia axyridis: An environmental risk assessment for northwest Europe. Biocontrol, 53:37–54.

van Lenteren, J., (2012). The state of commercial augmentative biological control: Plenty of natural enemies, but a frustrating lack of uptake. Biocontrol, 57.

Vorburger, C., Gehrer, L., and Rodriguez, P. (2010). A strain of the bacterial symbiont Regiella insecticola protects aphids against parasitoids. Biology letters, 109–111.

Vorburger, C., and Gouskov, A. (2011). Only helpful when required: a longevity cost of harbouring defensive symbionts. Journal of evolutionary biology, 24:1611–1617.

Walker, T., Johnson, P. H., Moreira, L. A., Moreira, L. A., Iturbe-Ormaetxe, I., Frentiu, F. D., McMeniman, C. J., McMeniman, C. J., Leong, Y. S., Dong, Y. D., Axford, J. K., Kriesner, P., Lloyd, A. L., Lloyd, A. L., Ritchie, S. A., O’Neill, S. L., O’Neill, S. L., and Hoffmann, A. A. (2011). The wmel Wolbachia strain blocks dengue and invades caged Aedes aegypti populations. Nature, 476:450–453.

Wilkinson, T., Fukatsu, T., and Ishikawa, H. (2003). Transmission of symbiotic bacteria buchnera to parthenogenetic embryos in the aphid Acyrthosiphon pisum (hemiptera: Aphidoidea). Arthropod structure development, 32(2):241–245.

Yang, Q., Gill, A., Robinson, K. L., Umina, P. A., Ross, P. A., Zhan, D., Brown, C., Bell, N., MacMahon, A., and Hoffmann, A. A. (2023). A diversity of endosymbionts across Australian aphids and their persistence in aphid cultures. Environmental microbiology, 25:1988–2001.

Yang, X., Pan, H., Yuan, L., and Zhou, X. (2018). Reference gene selection for rt-qpcr analysis in Harmonia axyridis, a global invasive lady beetle. Scientific reports, 8:2045–2322.

Zhang, D., Maiga, H., Li, Y. et al. (2024). Mating harassment may boost the effectiveness of the sterile insect technique for Aedes mosquitoes. Nature Communications, 15:2041–1723.

Zhao, Y., Nasrullah, Z., and Li, Z. (2019). Pyod: A python toolbox for scalable outlier detection. Journal of machine learning research, 20:1–7. employed in Python version 3.8.

Zytynska, E. S., Tighiouart, K., and Frago, E. (2021). Benefits and costs of hosting facultative symbionts in plant-sucking insects: A meta-analysis. Molecular ecology, 30:2483–2494.

