## Supplemental Table 1 for "Endosymbionts impact ladybird predation rates of aphids in a temperature-dependent manner"

### SUPPLEMENTAL MATERIAL

Supplemental Table 1: Log-rank (mantel-cox) Test testing difference between the species and clones of the aphids. Only the aphid groups without endosymbionts are investigated. The table denotes significance at the 0.05 level, denoted (*).

| **Uninfected ahids** | 14 ^◦^C | 20 ^◦^C | 26 ^◦^C |
| --- | --- | --- | --- |
| *R. padi* - : *M. persicae-A* *Rickettsiella* - | <0.0001 | <0.0001 | <0.0001 |
| *R. padi* - : *M. persicae*-B *Regiella* - | 0.0015 | 0.0073 | <0.0001 |
| *M. persicae-A* *Rickettsiella* - : *M. persicae*-B *Regiella* - | 0.0012 | <0.0001 | 0.0002 |
