## Supplemental Table 2 for "Endosymbionts impact ladybird predation rates of aphids in a temperature-dependent manner"

### SUPPLEMENTAL MATERIAL

Supplemental Table 2: Log-rank (Mantel-Cox) test results from the difference between the temperatures 14^◦^C, 20^◦^C and 26^◦^C. The table denotes significance at the 0.05 level (*). No significance is denoted NS. The table is divided according to species; large spotted ladybirds consuming *R. padi* (*Ri. viridis* infected, *Re. insecticola* infected and uninfected), two-spotted ladybirds consuming *M. persicae*, clone A, (*Ri. viridis* infected and uninfected), and two-spotted ladybirds consuming *M. persicae*, clone B, (*Re. insecticola* infected and uninfected).

| **Large spotted ladybird** | *R. padi* *Rickettsiella* + | *R. padi* - | *R. padi* *Regiella* + |
| --- | --- | --- | --- |
| 14 : 20 | <0.0001 | 0.0008 | <0.0001 |
| 20 : 26 | 0.0024 | <0.0001 | 0.0203 |
| 14 : 26 | <0.0001 | <0.0001 | <0.0001 |
| **Two-spotted ladybird** | *M. persicae*-A *Rickettsiella* + | *M. persicae-A* *Rickettsiella* - |  |
| 14 : 20 | <0.0001 | <0.0001 |  |
| 20 : 26 | <0.0001 | <0.0001 |  |
| 14 : 26 | <0.0001 | <0.0001 |  |
| **Two-spotted ladybird** |  | *M. persicae*-B *Regiella* - | *M. persicae*-B *Regiella* + |
| 14 : 20 |  | <0.0001 | 0.3193 |
| 20 : 26 |  | <0.0001 | <0.0001 |
| 14 : 26 |  | <0.0001 | <0.0001 |
